# TNF-α levels in respiratory samples are associated with SARS-CoV-2 infection

**DOI:** 10.1101/2021.07.12.452071

**Authors:** Matias J. Pereson, Maria Noel Badano, Natalia Aloisi, Roberto Chuit, MME de Bracco, Patricia Bare

**Affiliations:** Instituto de Medicina Experimental (IMEX) CONICET, Instituto de Investigaciones Hematologicas (IIHEMA), Academia Nacional de Medicina, Buenos Aires, Argentina; Universidad de Buenos Aires. Facultad de Farmacia y Bioquímica. Instituto de Investigaciones en Bacteriología y Virología Molecular (IBaViM). Buenos Aires, Argentina; Instituto de Investigaciones Epidemiológicas, Academia Nacional de Medicina

**Keywords:** SARS-COV-2, COVID-19, respiratory samples, Interleukin-6, TNF-α

## Abstract

**Purpose:** The aim of this study was to measure levels of IL-6 and TNF-α in respiratory samples from individuals with symptoms compatible with COVID-19 and analyze their association with SARS-CoV-2 presence.

**Methods:** SARS-CoV-2 detection was performed using the CDC (USA) real-time RT-PCR primers, probes and protocols. Cytokine concentrations were measured using commercial reagents based on enzyme linked immunosorbent assay (ELISA).

**Results:** TNF-α median levels were greater in COVID19 (+) symptomatic group (5.88 (1.36 - 172.1) pg/ml) compared to COVID19 (−) symptomatic individuals (2.87 (1.45 – 69.9) pg/ml) (p=0.0003). No significant differences were shown in IL-6 median values between COVID-19 (+) and (−) symptomatic patients (5.40 (1.7 - 467) pg/ml and 6.07 (1.57 – 466.6) pg/ml respectively). In addition, increased TNF-α levels (greater than 10 pg/ml), but not IL-6, were associated with SARS-CoV-2 presence (OR= 5.7; p=0.006; 95% CI= 1,551 to 19,11).

**Conclusions:** We found a statistically significant association between the production of local TNF-α and the presence of the virus in early stages of infection. IL-6 showed high levels in swabs from some symptomatic patients but independent from SARS-CoV-2 presence and viral load, individual’s age and gender. On the contrary, TNF-α evaluation confirmed the presence of inflammatory response but mostly related to COVID-19. More studies are required in order to characterize the cytokine profile expressed at the site of infection of SARS-CoV-2 and its implications in disease outcomes.

## Introduction

The coronavirus disease 2019 (COVID-19) caused by the severe acute respiratory syndrome coronavirus 2 (SARS-CoV-2) is one of the current major health concerns. After 15 months since the emergence of the pandemic SARS-CoV-2, up to 194 million cases were confirmed, including more than 4 million deaths worldwide due to severe forms of COVID-19 (WHO, 2021). Severe COVID-19 is widely considered an immunopathology disease driven by an excessive and deleterious immune response mounted against the pathogen rather than by the action of the pathogen *per se* (Mangalmurti & Hunter, 2020; Pedersen, & Ho, 2020; Hu et al. 2021). Many components of the innate immune system have been associated with this phenomenon and have been proposed as biomarker for COVID-19 progression. In this sense, high levels of serum interleukin-6 (IL-6) and tumor necrosis factor alpha (TNF-α) were correlated with increased morbidity and mortality in COVID-19 patients (Costella-Ruiz et al. 2020; Del Valle et al. 2020; Herold et al. 2020; Robinson et al. 2020a; Zhu et al. 2020; Danlos et al. 2021; Sabaka et al. 2021). However, innate immune mediators are necessary for efficient clearance of infectious agents. On the other hand, reports of lower median IL-6 levels in severe COVID-19 compared with other inflammatory conditions, such as acute respiratory distress syndrome and bacterial sepsis were published (Chen et al. 2020; Hambali et al. 2020; Chen & Quach, 2021). Additionally, many studies found no significant differences in TNF-α levels between severe and non-severe COVID-19 cases (Udomsinprasert et al. 2021). Therefore, the role of these cytokines in COVID-19 progression remains contradictory and needs further study.

To our knowledge, IL-6 and TNF-α concentration at the site of infection in COVID-19 patients has not been previously reported. Therefore, the aim of this study was to evaluate IL-6 and TNF-α concentration in swab samples from individuals showing symptoms compatible with COVID-19 who were positive or negative for SARS-CoV-2 genome detection.

## Material and methods

### Patients and study design

The study was conducted on 127 archived swab samples referred to the Academia Nacional de Medicina for diagnosis of SARS-CoV-2 by real-time polymerase reaction (RT-PCR) between August 2020 and April 2021. Samples were obtained during the initial stages of disease progression (1 to 10 days). Symptomatic individuals with detectable (n=52) or undetectable (n=33) SARS-CoV-2 infection were analyzed. A group consisting of asymptomatic individuals and negative for SARSCoV-2 genome (n=42) were included to obtain cytokines’ normal range.

### Collection of samples

Combined nasopharyngeal and oropharyngeal swabs were collected by trained healthcare staff, placed into a single tube pre-filled with 2 mL of saline solution (0,9% sodium chloride) and were transferred on the same day to the laboratory for SARS-CoV-2 genome detection. Samples were stored at −80 °C until cytokine assays were performed.

### SARS-CoV-2 genome detection and viral load

Genomic extraction was carried out with the QIAamp Viral RNA minikit (Qiagen) following manufacturer’s instructions. A real time-based methodology was performed for viral genome detection using the CDC RT-qPCR protocol for Wuhan virus (nCoV-19), 2019-nCov CDC USA. The procedure incorporates a set of oligonucleotides primers and double-marker hydrolysis probes (Taqman®) (2019 nCov_N1 and N2) that amplify regions of the viral nucleocapsid gene (N). An internal control for RNase P (RP) was also included. Viral load was calculated considering Ct values.

### IL-6 and TNF-α measurement

Concentrations of IL-6 (standard curve range: 0-300 pg/mL) and TNF-α (standard curve range: 0 −500 pg/mL) in swabs were determined using commercial reagents based on enzyme linked immunosorbent assay (ELISA) (BD-Biosciences, San Diego, California, United States). Recommendations of the supplier were followed for procedure. Duplicates were performed in selected samples to verify the accuracy of the results.

### Statistical analysis

Mann-Whitney U test was used for testing differences between two groups. Categorical variables were compared with Chi-square test or Fisher’s exact test. Confidence intervals were set at 95% (CI95). In all cases, a p value <0.05 was considered significant. Data and graphs were performed using the GraphPad Prism 9.1.0 software (GraphPad Software, San Diego, CA, USA).

### Ethics approval

Experimental protocols and procedures performed in this work have been approved by the Biosafety Review board of the IMEX-CONICET-Academia Nacional de Medicina and the Ethical Committee of the Academia Nacional de Medicina.

## Results

### Study population

Combined oro- and nasopharyngeal swabs from 127 patients submitted for diagnosis of COVID-19 were examined for SARS-CoV-2 genome presence and IL-6 and TNF-α concentration as described above. Since samples were drawn for SARS-CoV-2 infection diagnostic purposes, an important aspect of this study consisted in the timing of cytokine concentration assessment, performed during the first 10 days of symptoms in most of these patients. The most prevalent symptoms included malaise, myalgia, headache, and fever. Characteristics of the study population are shown in Table 1.

**Table 1.** Population characteristics

|  | COVID-19 (+)<br>Symptomatic pts | COVID-19 (-)<br>Symptomatic pts | COVID-19 (-)<br>Asymptomatic pts |
| --- | --- | --- | --- |
| Age (years) | 48,5 (0,16-93) | 56 (0,16-92) | 45 (0,7-93) |
| Gender (F/M) | 26/27 | 15/18 | 22/21 |
| Hospitalized pts (n) | 26 | 29 | 0 |
| SARS-CoV-2 viral load (Ct) | 29 (14-39) | na | na |
| Days of symptoms | 4 (1-11) | 2 (0-10) | na |

Values are expressed as median (range) unless otherwise noted. pts patients, *F* female, *M* male, *Ct* cycle threshold , *na* not applicable

### Cytokine analysis

Normal ranges of IL-6 and TNF-α for swab samples were set up using a group of asymptomatic COVID-19 (−) individuals. In this group, IL-6 median value was 3.18 pg/ml (0.78-8.39) and TNF-α median value was 4.72 pg/ml (3.19-9.69).

IL-6 was increased in both groups of symptomatic patients compared to asymptomatic patients (Fig. 1a). However, the difference between COVID-19 (+) and (−) symptomatic patients, with median values of 5.40 (1.7-467) pg/ml and 6.07 (1.57-466.6) pg/ml respectively, did not reach statistical significance (Fig. 1a). Despite both groups of symptomatic patients showed higher TNF-α levels compared to the asymptomatic group, no significant differences were observed (Fig. 1b). However, TNF-α levels were greater in COVID (+) symptomatic group [5.88 (1.36-172.1) pg/ml] compared to COVID (−) symptomatic individuals [2.87 (1.45–69.9) pg/ml], displaying a better relationship among this cytokine local concentration and the presence of SARS-CoV-2 infection (p=0.0003) (Fig. 1b). Considering a threshold value of 10 pg/ml for TNF-α, the presence of SARS-CoV-2 was associated with increased levels of TNF-α (OR= 5.7; p=0.006; 95% CI=1,551 to 19,11). On the contrary, IL-6 values did not show any association with the infection.

**Fig.1:**
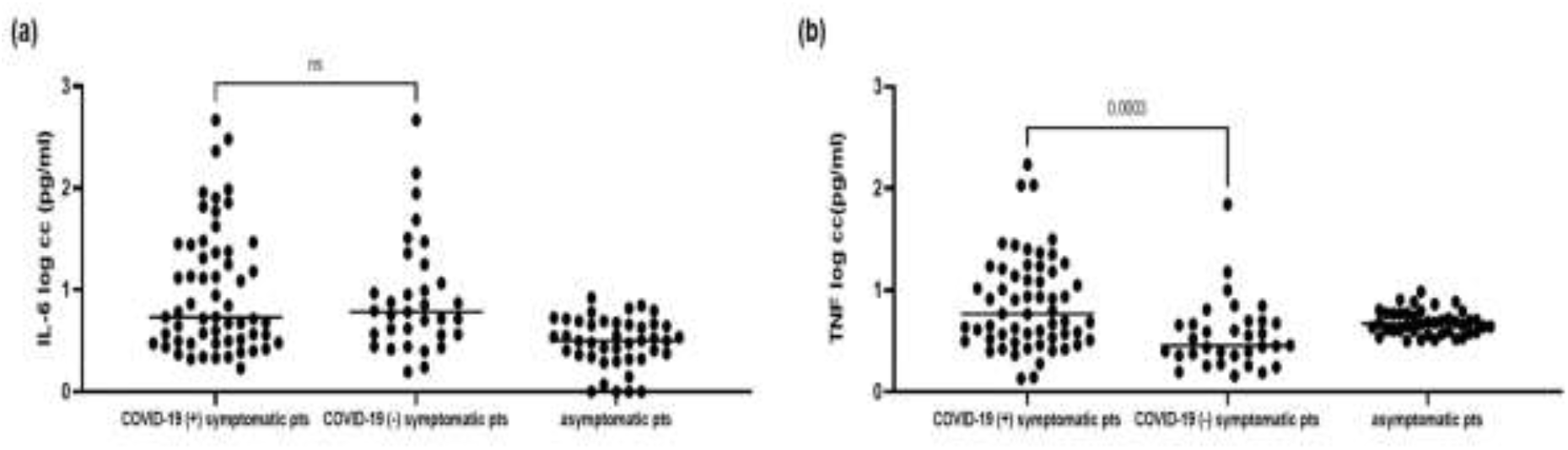
IL-6 (A) and TNF-α (B) concentrations (log pg/ml) in swab samples from symptomatic COVID-19 (+) and COVID-19 (−) symptomatic individuals.

Disease severity did not correlate with levels of IL-6 or TNF-α since median values in hospitalized and outpatients were not significantly different. However, COVID-19 (+) hospitalized individuals showed increased TNF-α values [median 4.89 pg/ml (1.4 – 172.2)] compared to COVID (−) hospitalized patients [median 3.36 pg/ml (1.4 – 70)] (p=0.0138). Among the group of COVID-19 (+) individuals, correlation between IL-6 or TNF-α levels and viral load (Ct) was not found. Furthermore, older age and gender did not seem to be related with higher values of IL-6 or TNF-α among the COVID-19 (+) group.

## Discussion

In the present study, we evaluated the concentration of IL6 and TNF at the entry site of SARS-CoV-2. Although levels of IL-6 and TNF-α were both increased in some patients during the first stages of COVID-19, only TNF-α was significantly associated with SARS-CoV-2 presence. This result may indicate a distinct cytokine profile expressed at this site during the symptomatic phase of SARS-CoV-2 infection.

Early in the pandemic, patients with high systemic levels of IL-6 and TNF-α were correlated with an excessive and deleterious immune response driving to COVID-19-related morbidity and mortality (Costella-Ruiz et al. 2020; Del Valle et al. 2020; Herold et al. 2020; Robinson et al. 2020a; Danlos et al. 2021; Sabaka et al. 2021). Focusing on IL-6 as a possible marker to predict the progress of COVID-19, several large studies suggested that systemic IL-6 levels of more than 80 pg/mL were the best laboratory predictor of respiratory failure and death and many IL-6 inhibitors therapies have been approved for the treatment of COVID-19 (Chen et al. 2020; Hambali et al. 2020; Chen & Quach, 2021; NIH, 2021; Udomsinprasert et al. 2021). However, many authors argued that although IL-6 levels may suggest severity of responses, they do not necessarily imply pathogenesis (Zhang et al. 2020; Chen & Quach, 2021; NIH, 2021; Udomsinprasert et al. 2021). Furthermore, some studies reported low median IL-6 levels in severe COVID-19 patients when compared to other inflammatory conditions, such as acute respiratory distress syndrome (ARDS) and bacterial sepsis (Calfee et al. 2014; Sinha et al. 2018). Accordingly, we could not find a correlation between IL-6 or TNF concentration in respiratory samples and severity of disease (analyzed as the requirement of hospitalization). In addition, IL-6 and TNF-α normal values were present even in severe hospitalized COVID-19 patients.

Likewise, reports on anti-TNF-α use are suggestive of a therapeutic benefit in patients with COVID-19 (Robinson et al. 2020a; Robinson et al. 2020b). Observational data from patients already on anti-TNF therapy show a reduced rate of COVID-19 poor outcomes and death compared with other immune-suppressing therapies (Gianfrancesco et al. 2020; Robinson et al. 2020a; Robinson et al. 2020b). Although randomized controlled trials are needed, these results may indicate that early anti-TNF therapy could be beneficial in a group of patients (Robinson et al. 2020b). In this study, we found a statistically significant association between the production of local TNF-α and the presence of the virus in early stages of infection. Therefore, measurement of local TNF-α, and probably some other mediators, could contribute to identify those individuals that may benefit from immunomodulatory and cytokine inhibiting therapies.

An increase of IL-6 and TNF-α and other cytokines in respiratory samples has been already described in numerous respiratory viral infections and has been associated with a worse prognosis of the disease (Seemungal et al. 2000; Patel et al. 2009; Ugonna et al. 2016; Vazquez et al. 2019). Our findings highlight the wide and pleiotropic role of IL-6 in local inflammation process and show that TNF-α estimation and its association to viral presence might be a valuable biomarker that could contribute to disease evolution prognosis. Through our results, we could hypothesize that a proportion of individuals might take advantage of the early and sequential measurement of cytokine concentration (at local and systemic levels) acting as supplementary biomarkers for COVID-19.

## Limitations

Because this study was done in samples obtained for COVID-19 diagnostic purposes, simultaneous serum samples were not drawn at the time of swab collection. Therefore, no comparison can be made with the systemic IL-6 or TNF-α values reported in other studies. Lack of monitoring cytokines at different time points after diagnosis is another limitation of our study.

## Declaration of Author Contributions

PB conceived the study; MNB and MJP designed the study protocol and carried out the clinical assessment; PB made the analysis and interpretation of the database. PB, MNB and MJP drafted the manuscript; MMEB and RC critically revised the manuscript for intellectual content. All authors read and approved the final manuscript.

## Acknowledgements

MJP was recipient of a Doctoral fellowship from the Consejo Nacional de Investigaciones Científico y Tecnológicas, CONICET. MNB was recipient of a fellowship from the National Academy of Medicine. PB is member of the National Research Council (CONICET) Research Career Program.

